# Reproductive mode, stem cells and regeneration in a freshwater cnidarian with post-reproductive senescence

**DOI:** 10.1101/203646

**Authors:** Flóra Sebestyén, Zoltán Barta, Jácint Tökölyi

## Abstract

In many basal metazoans both somatic and reproductive functions are performed by cellular derivatives of a single multipotent stem cell population. Reproduction can drain these stem cell pools, imposing a physiological cost with subsequent negative effects on somatic maintenance functions. In the freshwater cnidarian *Hydra oligactis* both asexual (budding) and sexual reproductive modes (production of resting eggs) are present, and both of these are dependent on a common pool of interstitial stem cells. Resting eggs tolerate abiotic conditions which neither the parental animals, nor asexual offspring can survive (e.g. freezing). Therefore, when facing unfavorable conditions and increased mortality risk, hydra polyps are expected to show higher differentiation of interstitial stem cells into germ cells (i.e. sexual reproduction), compared to other cell types needed for self-maintenance or asexual reproduction. Here, by comparing sexually and asexually reproducing individuals to non-reproductives, we studied the physiological costs of reproduction (size of interstitial stem cell pools, their somatic derivatives and regeneration rate, which is dependent on these cell types) in *H. oligactis* polyps from a free-living Hungarian population prior to the onset of winter. Sexual individuals (but not asexuals) were characterized by significantly smaller interstitial stem cell pools, fewer somatic derivatives (nematoblasts involved in food capture) and lower regeneration ability compared to non-reproductives. We also found a negative correlation between germ cell counts and stem cell numbers in males (but not in females). These results show that the physiological costs of reproduction are higher for sexual individuals. They also suggest that increased differentiation of stem cells into gametes might limit investment into somatic functions in hydra polyps. Exhaustion of cellular resources (stem cells) could be a major mechanism behind the extreme post-reproductive senescence observed in this species.

## Introduction

Sexual reproduction is ubiquitous in the natural world. Sex has clear evolutionary benefits over asexual reproduction (such as producing recombinant genotypes that have higher fitness under changing conditions), but it also entails costs (such as the cost of producing males; Maynard Smith 1978). Asexual reproduction does not entail these costs and can be evolutionarily favoured under special conditions (Crow 1994). Both sexual and asexual reproduction have a common cost: that of investing resources into offspring at the expense of self-maintenance of the parent (the physiological cost of reproduction; Calow 1979; Harshman & Zera 2007; Flatt & Heyland 2011). In animals, commonly studied costs of reproduction are the drain of specific macronutrients (e.g. amino acids, proteins or carbohydrates; (Zera and Zhao, 2006; Cotter et al., 2011)), micronutrients (e.g. dietary antioxidants; (Alonso-Alvarez et al., 2008; Catoni et al., 2008)) or metabolic reserves (e.g. body fat; Ellers, 1995). However, any factor in limited supply that is required by multiple life functions can mediate trade-offs between reproduction and somatic maintenance.

In animals with high tissue plasticity – like sponges, cnidarians and flatworms – stem cells might represent such a limiting factor. While in adult vertebrates stem cells have only limited plasticity (Weissman, 2000), in some invertebrates the adult body contains populations of highly flexible multi-or pluripotent stem cells (e.g. archeocytes in sponges, interstitial cells in some cnidarians, neoblasts in flatworms) which are responsible for the maintenance of a wide range of functions through their derivatives (Extavour and Akam, 2003; Juliano et al., 2010; Gold and Jacobs, 2013; Kumano, 2015). The strong role of these stem cells in self-maintenance is clearly seen in hydrozoans where, for instance, interstitial cells give rise to nerve cells, nematocytes (stinging cells usable once to capture food) and gland cells involved in digestion (Bode, 1996; Bosch, 2009; David, 2012; Plickert et al., 2012). The availability of these cells (i.e. cellular resources) is thought to determine growth rate and the magnitude of regenerative responses in sponges, corals and hydrozoans (Tardent, 1963;Lang da Silveira and Van’t Hof, 1977; Simpson, 1984; Rinkevich, 1996; Henry and Hart, 2005) and experimental elimination of stem cells in the freshwater cnidarian *Hydra* impairs several life functions related to the descendant cell types (Diehl and Burnett, 1964; Marcum and Campbell, 1978; Marcum and Campbell, 1978; Sugiyama and Wanek, 1993). On the other hand, multipotent interstitial cells are also strongly involved in reproduction: they are incorporated into asexual offspring during fission, fragmentation or budding (Simpson 1984; Bode 1996), produce resting bodies (gemmules and reduction bodies in sponges; Simpson 1984), and give rise to germ cells (Simpson 1984; Bosch & David 1987; Newark et al. 2008) or germline stem cells (a less potent stem cell lineage which can differentiate into gametes but not somatic cells; Nishimiya-Fujisawa and Kobayashi, 2012; Sato et al., 2006).

The common involvement of a single pool of multipotent progenitors in both somatic and reproductive functions, in theory, implies that increased investment into reproduction (either sexual or asexual) necessarily reduces differentiation of stem cells into somatic derivatives, thereby contributing to the physiological cost of reproduction (Rinkevich, 1996; Henry & Hart 2005). However, if the expected reproductive value of sexual and asexual offspring is not equal, then reproductive investment into these offspring types should also differ. Such a difference in reproductive value between offspring types could arise e.g. if expected survival rate of sexual and asexual offspring is not identical. Indeed, differential investment into sexual and asexual reproduction is commonly seen in several animal groups (e.g. ascidians: Yund et al. 1997; aphids: Nespolo et al. 2009; *Daphnia:* Innes & Singleton 2008; freshwater hydra: Kaliszewicz and Lipińska 2011). However, much less is known about the physiological consequences of this differential allocation.

The freshwater cnidarian *Hydra oligactis* is a species with a mostly temperate/arctic distribution in the Northern Hemisphere. *H. oligactis* polyps reproduce asexually throughout the year, but switch to sexual reproduction during the autumn (Schuchert, 2010). The most commonly invoked explanation for this switch in reproductive mode is to produce the resting eggs that can survive the winter (Reisa 1973). Based on laboratory experiments, sexual reproduction is followed by a senescence-like degeneration and increased mortality of polyps (Brien, 1953; Yoshida et al., 2006; Tomczyk et al., 2015; Tökölyi et al., in press; Schenkelaars et al., 2017). Post-reproductive degeneration is accompanied by marked changes in the cellular composition of hydra polyps: intersitital cell populations are strongly reduced while reproductive cells increase in number (Tardent, 1974; Yoshida et al., 2006). Because of these changes, post-reproductive senescence in hydra is hypothesized to be the consequence of “gametic crisis”, in which stem cell populations become exhausted due to excessive differentiation into reproductive cells, limiting their involvement in somatic functions (Brien, 1966; Tardent, 1968; Tardent, 1974; Bosch, 2009).

In this study, we investigated cellular composition and regeneration rate in *H. oligactis* polyps differing in reproductive modes (sexual, asexual and non-reproductive individuals), sampled from their natural environment during the autumn sexual period. Firstly, the role of cellular resources in mediating the trade-off between reproduction and self-maintenance is poorly understood in natural populations of any taxon. To date, the role of the stem cell pool in this trade-off is suspected mostly based on the negative linkage between traits depending on stem cells (like suppressed regeneration during reproduction (Campbell, 1967)), and the actual depletion of stem cells after initiation of sexual reproduction - representing a more direct role and limitation of these cells - has been reported only in a handful of cases (Littlefield, 1985; Yoshida et al., 2006; Gold and Jacobs, 2013). Post-reproductive senescence and stem-cell depletion has been described in *H. oligactis* in the laboratory (Yoshida et al., 2006), but little is known about this phenomenon under natural conditions. Furthermore, previous studies worked with a few laboratory strains of *H. oligactis* and it is unclear weather variation in reproductive strategies are associated with patterns of stem cell loss and changes in regeneration ability, as would be predicted from life history theory.

We hypothesized that reproductive value of sexual offspring should be higher because these can survive the winter, while asexual offspring cannot. As a consequence, sexual individuals should invest more into reproduction, which would result in higher overall physiological cost of reproduction. Supporting the mediator role of the stem cell pool, this would manifest itself in lower availability of stem cells, their somatic derivatives and somatic functions depending on stem cells as well. Accordingly, we predicted lower number of stem cells, fewer nematoblasts (indicating reduced differentiation into somatic functions) and lower ability to regenerate in sexual individuals compared to asexuals or non-reproductives. Furthermore, we also predicted that, if differentiation of stem cells into reproductive function is traded off with somatic maintenance, then the number of reproductive cells should be negatively related to interstitial stem cells and their somatic derivatives.

## Materials & methods

### Collection of animals and culture conditions

Experimental animals were collected from an oxbow lake near Tiszadorogma in Eastern Hungary (47.6712N, 20.8641E). To determine the reproductive status of the animals in the lake and hence to detect the start of the autumn reproductive period we visited the site on 2^nd^ and 16^th^ October 2016 and collected N = 168 (N = Number of collected animals) and N = 136 hydra polyps, respectively. Further collections were performed four times in 2016: 26^th^ October (N = 127), 2^th^ November (N = 332), 15^th^ November (N = 121) and 6^th^ December (N = 51). Animals collected on the first two dates were not used in regeneration experiments or for quantification of cellular composition, only for the detection of sexual reproduction period. We collected animals from several sites along the shoreline of the lake to reduce the chance of obtaining genetically identical clones produced by asexual budding. Hydras were picked up from submerged vegetation, placed in Eppendorf tubes and brought to the laboratory in a cool box on the same day. In the laboratory, we recorded mode of reproduction according to three categories: (1) no reproduction (polyps without buds or gonads, N=272), (2) asexual reproduction (polyps with at least one bud, N=204), or (3) sexual reproduction (polyps with differentiated or developing gonads; this latter was defined as a thick, opaque swelling around the gastric region of the body column, N=155). Sexual individuals were further divided into three categories: males (polyps with differentiated testes, N=25), females (polyps with differentiated eggs, N=34), and sexual individuals in which sex could not be determined (N=96). This latter category included immature males and females with developing testes or eggs, and post-reproductives showing the morphological characteristics of sexual reproduction, but without clearly defined reproductive organs, since these categories are not unambiguously distinguishable. Animals were clearly referable to only one category of reproduction modes (we found just two asexual animals showing the morphological sign of sexual reproduction; these were coded as sexual individuals because gonadogenesis was clearly initiated).

### Head regeneration measurements

About half of the collected animals (altogether 338) were randomly assigned to head regeneration measurements, which were initiated one day after each of the four collections. Animals were decapitated below the tentacles, which means that the removed part contained the oral tip (i.e. the hypostome), the tentacles and a short part of the trunk (~10% of the body length). During regeneration we kept the animals individually in 24-well plates in ~ 3ml standard hydra medium (1.0 mM CaCl_2_, 0.1 mM MgCl_2_, 0.03 mM KNO_3_, 0.5 mM NaHCO_3_ 0.08 mM MgSO_4_; Zhang et al., 2002). We placed the plates with hydras in a Memmert ICP 700 climate chamber and kept them on constant photoperiod (16 h dark/ 8 h light cycle) and temperature in accordance with natural habitat temperature measurements on the four consecutive dates (12 °C, 9 °C, 5 °C, 4 °C, measured approximately 20 cm below the water surface on the day of collection). Hydras completely regenerate their head after 48-72 h on 18 °C (Ambrosone et al., 2012), but at lower temperature cell cycle and cell division is slower (Begasse et al., 2015), thus regeneration takes longer (Lillie and Knowlton, 1897). For this reason, we recorded regeneration 4 days after decapitation by a binary code system, based on the presence or absence of newly emerged tentacles.

The hypostome and tentacles amputated for the head regeneration experiments were used for species determination. *H. oligactis* can be distinguished from other *Hydra* species occurring at this site based on nematocyte morphology, which can be observed under a light microscope.

### Cell number measurement

One day after collection, we randomly selected a subset (altogether 155 animals) from the remainder of the animals for cell number measurement. These were macerated according to the standard procedure described by David (1973), and then cells were spread on a microscopic slide. Sample size for the cell number measurement was determined by time constraints: we only used samples for which macerations could be prepared on the next day after collection, such that cellular composition measured by us is as close as possible to the condition of animals at the time of sampling. Cellular composition was quantified within a few days after maceration. For each sample we recorded the number of epithelial cells, interstitial stem cells (large single interstitial cells or nests of two interstitial cells were recorded together to obtain an estimate of the frequency of stem cells; (Bosch and David, 1987), nematoblast nests (total number of nests of 1, 2, 3-4, 5-8 or >8 cells) and reproductive cells (sperm/sperm precursor nests in males and nurse cells in females). Reproductive cells at later stages of development are distinguishable from interstitial cells based on morphological criteria, as follows. In males, interstitial cell nests commited to sperm development increase in number and size and flagella start to develop on the sperm precursors (Littlefield, 1985). We used the presence of flagella as a morphological criterion to identify sperm cells/sperm precursors and counted the number of sperm/sperm precursor nests (i.e. groups of sperm precursors) with flagella to obtain a semiquantitative estimate of germ cell numbers in males (in this estimate all sperm precursor nests were pooled irrespective of the number of cells in them; this was necessary because the large number of individual germ cells in some nests made exact counting of cell numbers impractical). In females, interstitial cells commited to germ cell differentiation first increase in size and develop into nurse cells, which can be distinguished from interstitial cells by their larger cytoplasm volume (Zihler, 1972). Cells were identified as nurse cells when the diameter of the nucleus was equal or less than half of the cell diameter, indicating relatively large cytoplasm volume. This corresponds to Stage B oogonia in Zihler's (1972) notation. Only a small subset of these nurse cell develop into oocytes, but these incorporate neighbouring nurse cells through phagocytosis (Miller et al., 2000). Hence all nurse cells contribute to reproduction and therefore we counted all of them.

In all samples we systematically traversed slides until at least one hundred epithelial cells were recorded, and noted any other cell types alongside these epithelial cells. The median number of cells/cell nests recorded per sample (including the epithelial cells) was 208 (range: 107-3048).

The head region of the animals assigned to investigation of cellular composition was removed in the same way as in the head regeneration experiments and used for species determination. All sexual individuals involved in cell number measurements were categorized as males or females, based on the presence of mature gonads and / or sperm cells / nurse cells in macerates.

### Statistical analysis

The effect of reproductive mode on head regeneration was analyzed using Generalized Linear Mixed Models (GLMM) with binomial distribution. Our model contained regeneration (presence or absence) as dependent variable and reproduction mode as predictor. We included collection date as a fixed effect, to control for seasonal and temperature differences. We included collection site as a random effect to control for the possibility that animals from the same sampling point might be more similar to each other than to individuals from other sites because of shared environment or because some of them might be asexual descendants of a single individual. Binomial GLMMs were implemented in a Bayesian framework, employing the MCMCglmm R package (Hadfield, 2010; R Core Team, 2017). A Bayesian approach was required because our data suffered from complete separation (some experimental groups contained only non-regenerating animals). This problem can be circumvented in a Bayesian setting by setting a weak prior on fixed effects in MCMCglmm. We ran this model two times for our data sets then averaged the two results.

For testing the effect of reproductive mode on nematoblast and interstitial cell number, we used Poisson GLMM also implemented in a Bayesian framework. We included epithelial cell number as a fixed effect, because the number of epithelial cells was not exactly identical (sometimes we counted slightly more than one hundred); by controlling for epithelial cell number we take into account variation in stem cell numbers arising from slightly unequal sampling. We also included collection date as a fixed effect and collection site as a random effect for the reasons mentioned above. For analyzing the relation between sperm/nurse cell number and interstitial or nematoblast cell number (all cell type numbers were normalized to epithelial cell number), we performed Spearman rank correlation. All analyzes were performed in the R Statistical Environment (R Core Team, 2017).

## Results

### Reproductive phenology

None of the individuals collected on 2^nd^ October showed signs of sexual development. A single male bearing mature testes was observed on 16^th^ October. The proportion of sexual individuals on subsequent dates was 20.5%, 29.8%, 22.3% and 5.9% on 26^th^ October, 2^th^ November, 15^th^ November and 6^th^ December, respectively (Fig. 1.). The proportion of asexual animals was 18.1%, 30.1%, 46.3% and 49%, on the respective dates (Fig. 1.).

**Fig. 1.**
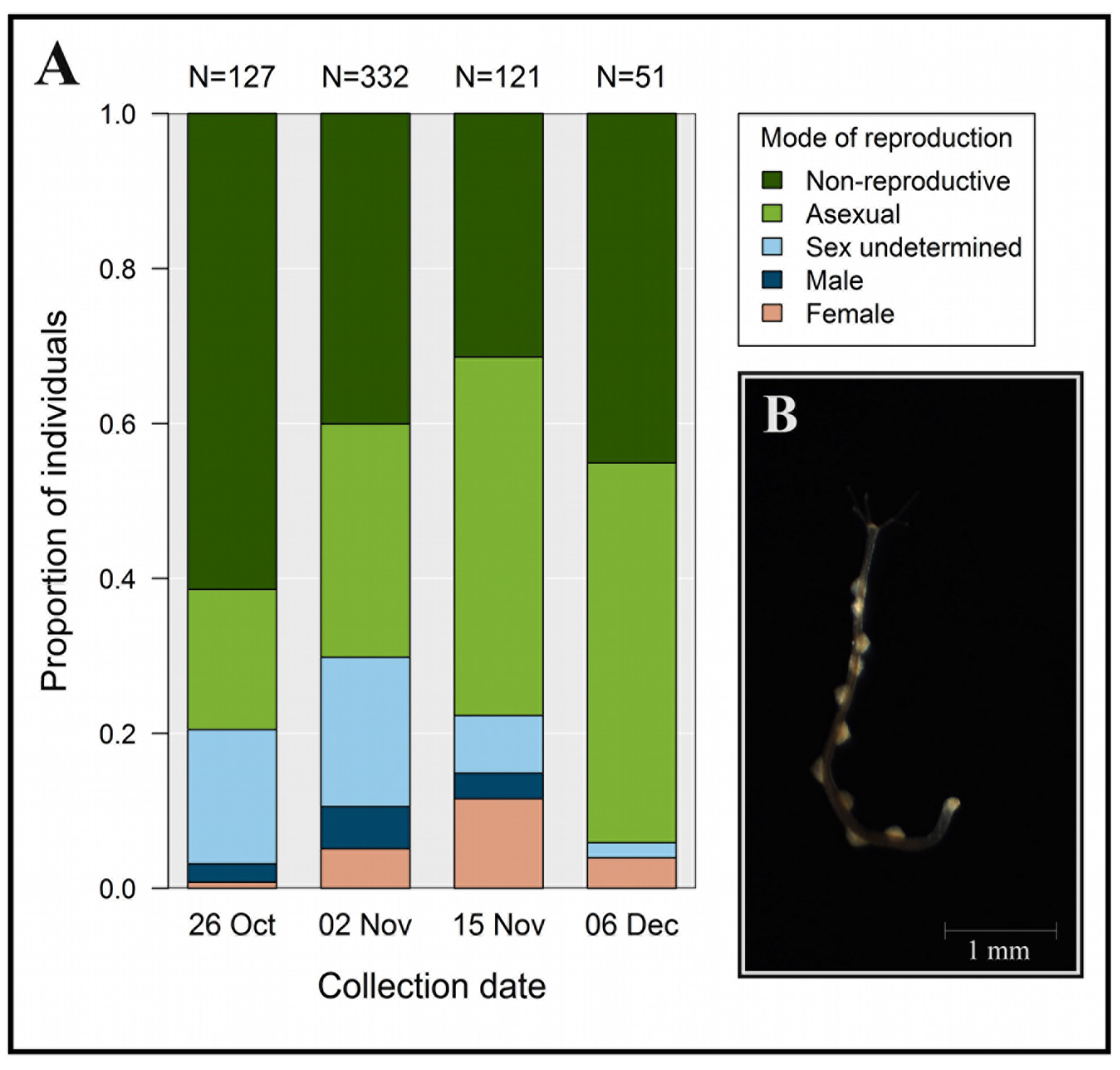
Proportion of individuals in different reproduction mode categories on four collection dates. Total sample sizes are shown above the bars (A). Reproductive mode categories were: non-reproductive (polyps without buds or gonads), asexual (polyps with at least one bud) and sexual (polyps with differentiated eggs (females), testes (males) or developing gonads (sex undetermined)). (B) Photograph of a wild-collected male polyp showing signs of post-reproductive degeneration (depleted testes and strongly reduced tentacles), collected on 02 Nov.

### Head regeneration

Regeneration abilities differed between reproduction mode categories (Fig. 2). Compared to non-reproductive animals (56.85% regenerated heads within 4 days), the proportion of animals regenerating heads was significantly lower in males (0%; posterior mean = −5.431, lower 95% CI = −8.855, upper 95% CI = −2.157, p<0.001), females (0%; posterior mean = −4.667, lower 95% CI = −8.545, upper 95% CI = −1.542, p<0.001) and animals with undetermined sex (23.21%; posterior meaN = −2.341, lower 95% CI= −3.353, upper 95% CI= −1.351, p<0.001). In asexual hydras, head regeneration did not differ significantly from non-reproductive individuals (30.61%; posterior mean = 0.568, lower 95% CI = −1.447, upper 95% CI = 0.252, p = 0.179). Collection date as a fix effect had significant effect on regeneration rate: compared to the first collection, regeneration rate was significantly lower in all dates (results not shown).

**Fig. 2.**
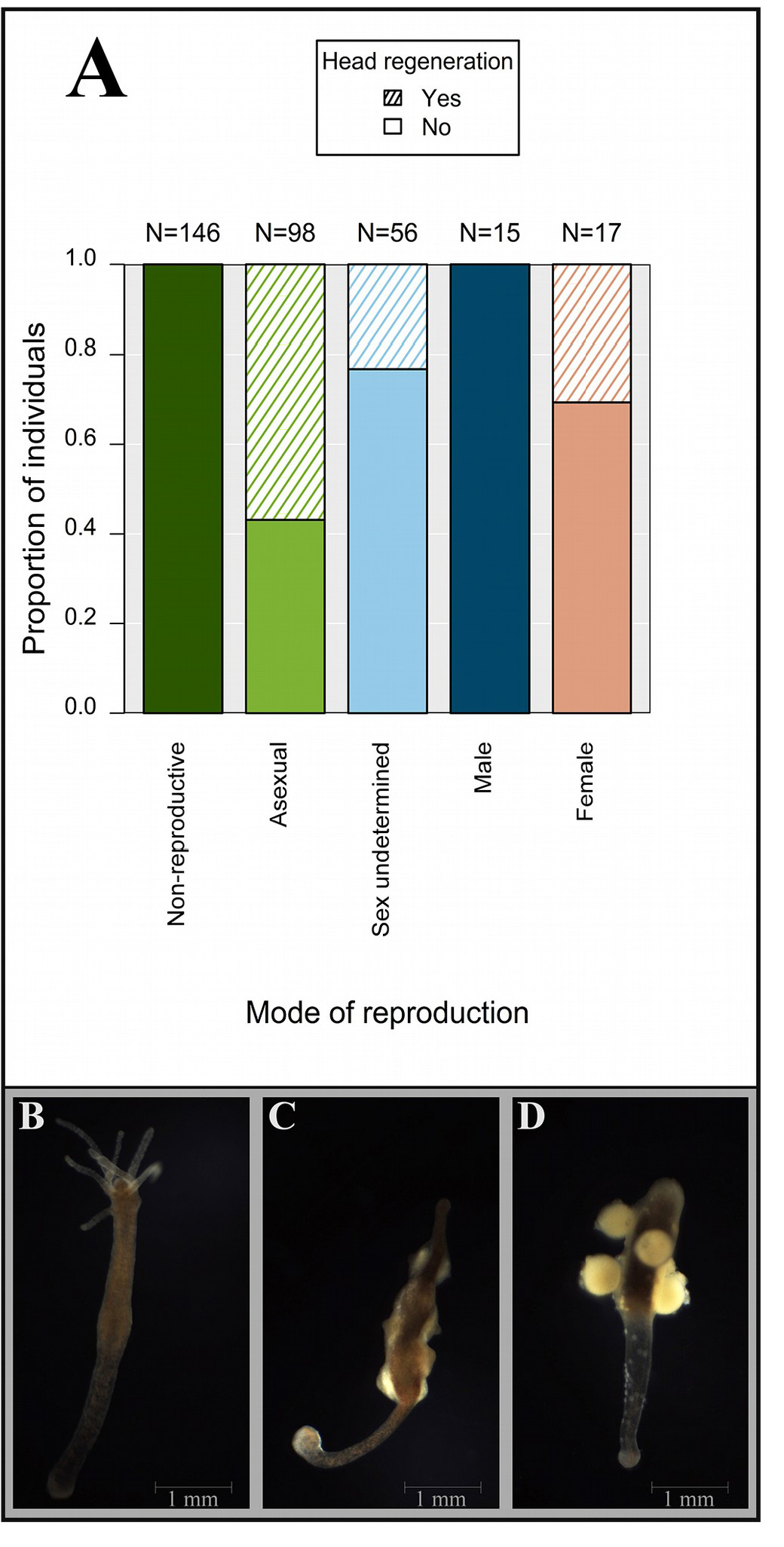
Head regeneration (presence or absence of tentacles) 4 days after decapitation in hydras differing in reproductive mode (A). Non-reproductive (B), male (C) and female (D) polyp after decapitation illustrating the markedly reduced head regeneration ability of sexual individuals. See Fig. 1. for reproductive mode categories. Photographs were taken after finalization of regeneration experiments (8 days post-amputation).

### Nematoblast and interstitial cell number and mode of reproduction

Mode of reproduction had a significant effect on both nematoblast cell number and interstitial cell number (Fig. 3). Compared to non-reproductive animals, interstitial cell number did not differ in asexually reproducing individuals (posterior mean = 0.195, lower 95% CI = −0.331, upper 95% CI = 0.752, p = 0.494), but it was lower in males (posterior meaN = −0.923, lower 95% CI = −1.602, upper 95% CI = −0.23, p=0.008) and females (posterior mean = −1.254, lower 95% CI = −1.753, upper 95% CI = −0.792, p < 0.001). Nematoblast cell number was significantly lower in males (posterior mean = −1.196, lower 95% CI = −1.937, upper 95% CI = −0.416, p < 0.001) and females (posterior mean = −1.929, lower 95% CI = −2.488, upper 95% CI = −1.383, p < 0.001), but it was marginally significantly higher in asexual animals (posterior mean = 0.578, lower 95% CI = −0.023 upper 95% CI = 1.144, p = 0.055), compared to non-reproductives.

**Fig. 3.**
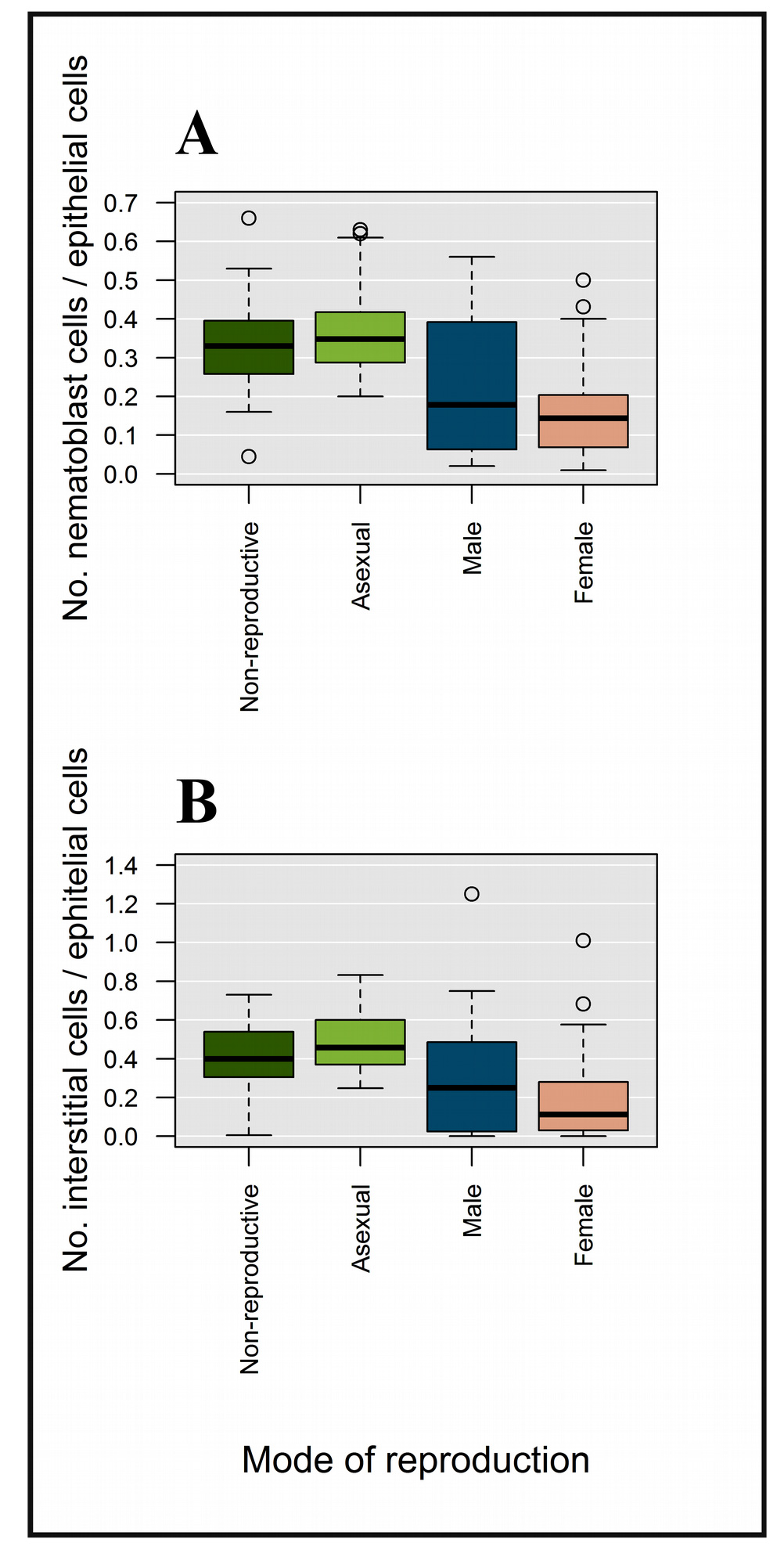
Nematoblast number (A) and interstitial stem cell number (B) of individuals in different reproductive mode categories. All sexual individuals were categorized as males or females based on the presence of mature gonads and / or sperm cells / nurse cells in macerates.

### Gamete and interstitial cell number

There was a significant negative correlation between number of sperm precursor nests and interstitial cell number (Spearman correlation, ρ= −0.764, p<0.001, N=22), as well as number of sperm precursor nests and nematoblast cell number in males (Spearman correlation, ρ=−0.852, p<0.001, N=22) (Fig. 4). We found a significant positive correlation between nurse cell number and interstitial cell number in females (Spearman correlation, ρ=0.424, p=0.002, N=51), but there was no correlation between their nurse cell number and nematoblast number (Spearman correlation, ρ=0.028, p=0.844, N=51) (Fig. 4). There was a significant positive correlation between nematoblast and interstitial cell counts in both males (Spearman correlation, ρ= 0.578, p=0.008, N=22) and females (Spearman correlation, ρ=0.448, p=0.001, N=51).

**Fig. 4.**
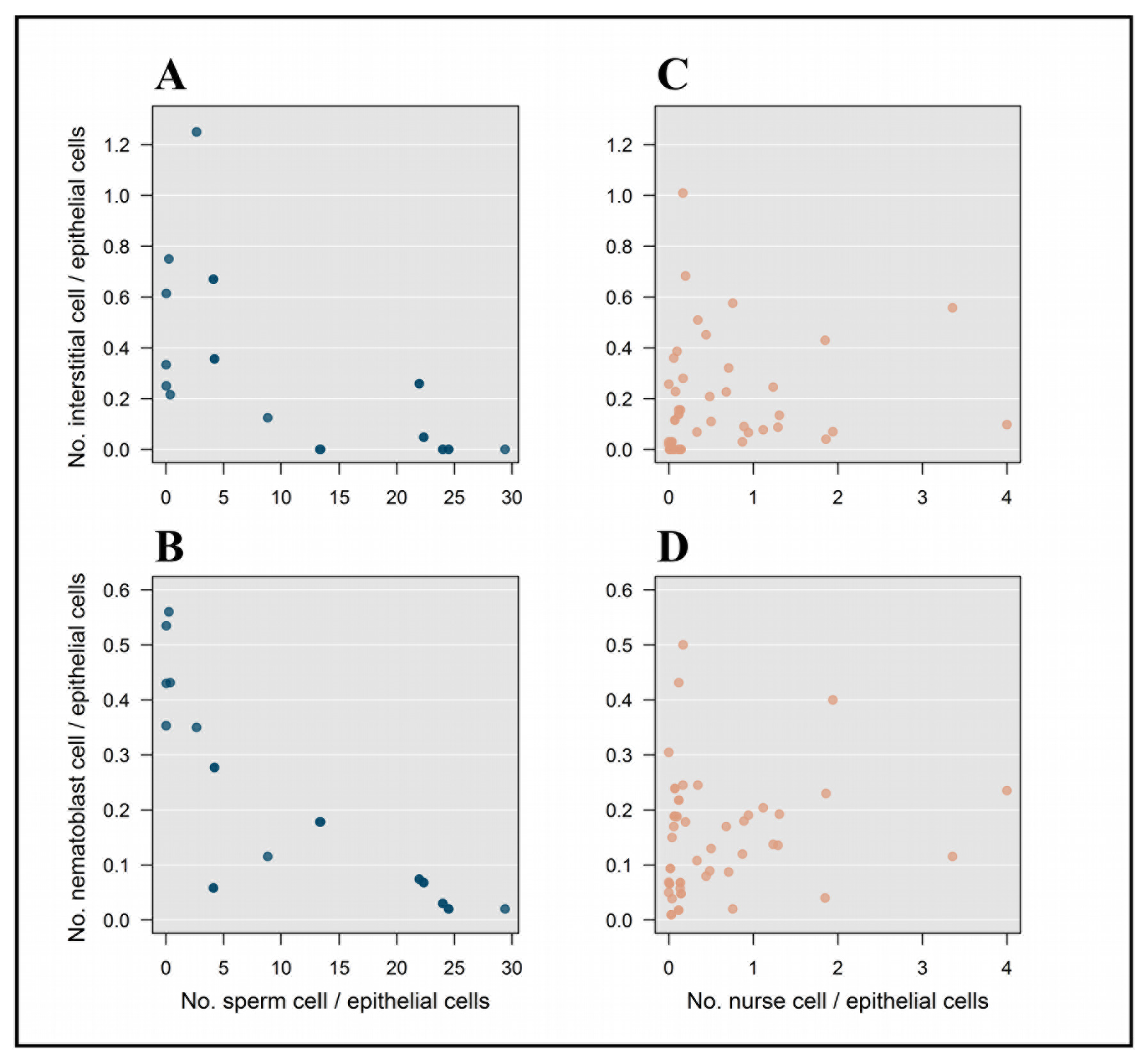
Correlation between reproductive cell number (sperm or nurse cells) and interstitial and nematoblast cell number in females (A and B) and interstitial and nematoblast cell number in males (C and D). All cell numbers were normalized to epithelial cell number.

## Discussion

In this study we described regeneration rate and cellular composition of *H. oligactis* polyps differing in reproductive strategies. We found that sexual (but not asexual) reproduction is associated with reduced regeneration ability and decreased number of interstitial stem cells and nematoblasts involved in food capture. This observation indicates that sexual reproduction is associated with increased physiological cost. Our results lend support to the hypothesis that life history decisions in *Hydra* might be mediated by a competition for a limited stem cell pool involved in multiple life functions (Rinkevich, 1996).

In animals with high tissue plasticity, the activity of multipotent stem cells is required for multiple life functions. Reduced availability of stem cells has been suggested to be involved in the determination of *Tubularia* hydrant lifespan (Tardent, 1963), and also thought to be responsible for post-reproductive degeneration in *H. oligactis* (Brien 1966; Tardent, 1968; Tardent, 1974; Bosch, 2009). Increased commitment of stem cells into germ cells likely reduce the differentiation of stem cells into somatic cells (a process termed "gametic crisis"; (Brien, 1966; Bosch, 2009), possibly causing a decline in survival of *H. oligactis* (Yoshida et al., 2006). Our observation that interstitial cells and nematoblasts were reduced, while germ cell numbers increased during sexual reproduction in animals from a natural population are in accordance with findings obtained under laboratory circumstances in this species (Yoshida et al., 2006). Although other *Hydra* species do not seem to show similar patterns of senescence, there is evidence that similar exhaustion might occur in *Aurelia* polyps that have been stimulated to strobilate (generate sexual medusa) many times (Gold and Jacobs, 2013).

In parallel to the decline in stem cell pools, regeneration rate was also reduced in sexual polyps. Regeneration is a somatic function that likely depends on the availability of cellular resources (stem cells). Stem cells are crucial in all types of regeneration either because they proliferate to produce cells that will be involved in regeneration or because they migrate to the wound site to re-form lost body parts (Sánchez Alvarado, 2000; Bely and Nyberg, 2010; Sugimoto et al., 2011). Any physiological process that reduces the availability of stem cells can therefore, in theory, limit regeneration (Kramarsky-Winter and Loya, 2000; Henry and Hart, 2005). Indeed, reduced availability of cellular resources has been invoked previously to explain the reduction in regenerative responses in response to subsequent amputations (Gross, 1925; Kanajew, 1926; Tardent and Tardent, 1956; Tardent, 1963) and the suppression of regeneration after sexual reproduction (Campbell, 1967 or vice versa: Rinkevich and Loyla, 1989).

The trade-off between differentiation into somatic and reproductive functions is further underscored by our observation that interstitial stem cell numbers and nematoblast numbers were negatively related to germ cell counts in males. Such a negative relationship could arise because individuals with a higher reproductive investment (larger germ cell counts) have fewer remaining interstitial cells / nematoblasts. Interestingly, we observed no relationship between reproductive cell numbers and interstitial cell / nematoblast counts in females. This could mean that the trade-off between germ cells and somatic cell types is different in females and males. However, it might also have been caused by differences in the way reproductive investment is estimated from reproductive cell counts. Specifically, in females nurse cells become incorporated into developing eggs, and during egg maturation fewer and fewer nurse cell will be located separately (Zihler, 1972; Miller et al., 2000). As a consequence, females would show a progressive reduction in both nurse cell numbers and depletion of interstitial stem cells / nematoblasts during egg maturation if stem cells differentiate into nurse cells and these become incorporated into eggs. This could explain the relatively high number of females in which nurse cells, interstitial cells and nematoblasts were all depleted (Fig. 4).

In spite of the strongly reduced stem cell numbers in sexually reproducing polyps (which suggest that interstitial cells are converted to germ cells during gonadogenesis), the exact explanation for interstitial cell depletion during sexual reproduction in *H. oligactis* is still not clear. Current models of germ cell specification suggest that there are two interstitial stem cell lineages in *Hydra*, which are morphologically indistinguishable (reviewed in Nishimiya-Fujisawa and Kobayashi, 2012). Multipotent stem cells (MPSCs) give rise to somatic cells, like nerve cells, gland cells and nematocytes, while germline stem cells (GSCs) are unable to produce somatic cells but differentiate into nurse cells and sperm. GSCs derive from MPSCs (Nishimiya-Fujisawa and Kobayashi, 2012), hence MPSCs are able to produce both somatic and reproductive cells. This is supported by observations that (1) *H. oligactis* polyps in which GSCs were experimentally ablated are still able to quickly develop germ cells when exposed to cold temperature (Littlefield et al., 1985) and (2) cloning individual interstitial cells in another *Hydra* species *(H. magnipapillata)* can give rise to both somatic and germline cells (Bosch and David, 1987). Because of the complexity of interstitial stem cell lineages in hydra, the gaps in knowledge of their dynamics and the indistinguishability of the two major stem cell types, it is possible that only a subset of the cells identified in this study as interstitial cells (MPSCs) take directly part in the somatic-reproductive trade-off. However, since these MPSCs can differentiate into GSCs (or directly into germ cells), the trade-off in stem cell differentiation between somatic and reproductive functions remains the same. Future studies of stem cell differentiation during gonadogenesis and hydra germline stem cells would help to elucidate the exact mechanisms behind stem cell depletion during gametogenesis observed in this species.

While sexual reproduction was associated with reduced somatic cell types and regeneration ability, we did not observe such a reduction in asexual individuals. Interestingly, this pattern mirrors the phylogenetic distribution of reproductive mode and regeneration ability in several invertebrate groups: regenerative capacities are lower in sexual species in segmented worms (Zattara and Bely, 2016) and flatworms (Peter et al. 2001), compared to asexual ones. The higher stem cell pools and regenerative potential of asexual polyps clearly indicates that this type of reproduction does not impose such a high physiological cost on the parent polyp as sexual reproduction does. Indeed, asexual buds in *Hydra* are thought to be produced from excess cells arising from an actively dividing stem cell population (Bosch, 2009; Gold & Jacobs, 2013), in which case they are less likely to drain from the limited resources of the parent. However, since asexual buds are prone to freezing just as the parent animal, they are likely to have a lower reproductive value during autumn than resting eggs. Hence, asexuals in this population appear to follow a strategy of producing offspring with low reproductive value at a low cost, as opposed to sexuals, which produce offspring of high reproductive value at a high physiological cost. This latter strategy might be considered a case of terminal investment (Williams, 1966; Clutton-Brock, 1984).

In addition to describing patterns of reproductive mode, stem cells and regeneration in *H. oligactis*, we also provide data on the natural phenology of sexual reproduction for this species. While gametogenesis in *H. oligactis* is known to occur during the autumn and to last until early winter, the ecology of *H. oligactis* has been investigated by only a handful of studies so far (Welch and Loomis, 1924; Miller, 1936; Bryden, 1952; Ribi et al., 1985) Sexual reproduction is thought to occur in this species when adult survival is expected to be low due the cold temperature and high risk of freezing (Reisa, 1973). However, previous studies have shown that, in general, only a subset of the population reproduces sexually at any time; moreover, sexually reproducing animals are not found in some years (Miller, 1936; Bryden, 1952; Ribi et al., 1985). In this Hungarian population, the proportion of sexually reproducing individuals was also lower than that of agonadic individuals (Fig. 1). Interestingly, the proportion of asexual animals showed an increasing tendency towards the onset of winter (even though the temperature was decreasing), possibly because sexual individuals were disappearing or reverted back to asexual reproduction (which is known to occur in individuals collected from this population under laboratory conditions; (Tökölyi et al., in press). Together with the observations that (1) initiation of sexual reproduction in *H. oligactis* strongly depends on the rate of the temperature drop (Kaliszewicz, 2015) and (2) some *H. oligactis* strains seem to have a lower propensity to initiate sexual reproduction (Tökölyi et al., in press), these results suggest that sexual reproduction in *H. oligactis* is a conditional and polymorphic strategy or maybe a form of bet-hedging (a stochastic switching between phenotypic states − a way of adaptation to fluctuating environment, e.g. (Cohen, 1966)), possibly determined by specific environmental conditions of the natural habitat (e.g. the risk of freezing).

Overall, our results suggest that sexual reproduction imposes a high physiological cost on *Hydra oligactis* polyps. The reduced regeneration abilities and depletion of stem cells in sexually reproducing animals compared to non-reproductives might imply that current sexual reproduction is an irreversibly induced reproduction strategy, and gamete production is prioritized over the maintenance of somatic functions and future survival during autumn. The highly divergent life history decisions of *H. oligactis* provide a great model system to study aging and non-senescent life history tactics and its physiology within a single species. In addition, in order to clarify the role of limiting cellular factors, further studies focusing on common cellular pools required by life history traits are much needed.

## Acknowledgements

This study was supported by NKFIH grant FK 124164. JT was supported by the ÚNKP-17-4 New National Excellence Program of the Hungarian Ministry of Human Capacities. ZB was supported by the NKFIH grant K 112527. We acknowledge the financial support of this work by the Hungarian State and the European Union under the EFOP-3.6.1 project.

